# Quantitative Personality Predictions from a Brief EEG Recording

**DOI:** 10.1101/686907

**Authors:** Wenyu Li, Chengpeng Wu, Xin Hu, Jingjing Chen, Shimin Fu, Fei Wang, Dan Zhang

## Abstract

The assessment of personality is crucial not only for scientific inquiries but also for real-world applications such as personnel selection. However, most existing ways to quantify personality traits rely on self-reported scales, which are susceptible to biases such as self-presentational concerns. In this study, we propose and evaluate a novel implicit measure of personality that uses machine learning (ML) algorithms to predict an individual’s levels in the Big Five personality traits from 5 minutes of electroencephalography (EEG) recordings. Results from a large test sample of 196 volunteers indicated that the personality scores derived from the proposed measure converged significantly with a commonly used questionnaire, predicted behavioral indices and psychological adjustment in a manner similar to self-reported scores, and were relatively stable across time. These evaluations suggest that the proposed measure can serve as a viable alternative to conventional personality questionnaires in practice.

## Introduction

Over a hundred years of scientific inquiry into individual differences has identified five overarching traits as the fundamental dimensions of personality: extraversion, neuroticism, conscientiousness, agreeableness, and openness to experience(McCrae & Costa, 2008; McCrae & John, 1992). These “Big Five” traits represent dispositional differences in cognitive, affective, behavioral and motivational patterns, and can predict important life outcomes such as psychological adjustment(Ozer & Benet-Martinez, 2006). Given the importance of the Big Five traits, it is crucial to develop a reliable measurement of them not only for academic research, but also for application scenarios such as personnel selection.

Most applications of the Big Five model rely on self-reported scales which require the respondents to read statements or adjectives which they judge in relation to their personality and report their degree of agreement(Costa Jr & McCrae, 2008; Gosling, Rentfrow, & Swann, 2003). These self-reported scales, whilst having the advantages of straightforwardness and cost-effectiveness, are susceptible to biases such as social desirability or self-presentational concerns. For example, a job applicant may deliberately fake his/her responses to a personality questionnaire to show competency for the position. This disadvantage limits the method’s effectiveness in certain application settings.

One way to tackle this problem is to use indirect measures that do not require the participants to report a subjective assessment of their own personality but make inferences from other sources of data such as observed behavioral patterns(Gawronski & De Houwer, 2014). Throughout the history of personality science, there have been multiple attempts to develop such measures. For example, psychoanalysts have used the subjective interpretation of ambiguous inkblot patterns to probe one’s unconscious mind(E. Exner Jr, 2003; Rorschach, 1921). However, its validity has been an ongoing issue of debate(Wood & Lilienfeld, 1999). A more recent example is the personality measure based on the Implicit Association Test (IAT), which employs measures of reaction time to assess the association strength between one’s concept of self and the concept of a trait(Schmukle & Egloff, 2005; Schnabel, Asendorpf, & Greenwald, 2008). These IAT-based measures have been demonstrated to have adequate reliability and validity, although what they actually measure may be conceptually distinct from explicit measures of personality(Dentale, Vecchione, & Barbaranelli, 2016).

In recent years, the introduction of machine learning techniques into psychological science has opened up new possibilities for implicit personality measures(Bleidorn & Hopwood, 2018). The machine learning approach to personality assessment focuses on developing automated algorithms to predict one’s personality from certain data sources, and the algorithms are usually cross-validated to ensure their generality to new samples. Recently, there have been reports of success in the application of this approach on individual’s digital footprints on social media websites(Settanni, Azucar, & Marengo, 2018; Wald, Khoshgoftaar, & Sumner, 2012; Wu, Kosinski, & Stillwell, 2015). For example, Wu et al.(Wu et al., 2015) developed machine learning models to predict one’s levels on the Big Five traits from Facebook “Likes”. The accuracy of their model’s predictions, evaluated against self-reported personality scores and predictive validity for life outcome variables, was higher than the judgments made by human informants.

Besides online behaviors, another type of data that may benefit from a machine learning approach is neurophysiological data. It has been an ongoing endeavor for psychologists and neuroscientists to investigate the neurobiological basis of personality(R. Jiang et al., 2018; Korjus et al., 2015; Nostro et al., 2018). Despite the fact that consensus has not been reached for many traits, broadly speaking, the available data do suggest that there are stable patterns of intraindividual variance in neural activities which correspond to dispositional differences at the behavioral level. However, for the purpose of developing neural-based personality measures, the existing studies are limited in two ways. First, many of the findings were obtained by techniques such as functional magnetic resonance imaging (fMRI), which due to their expensive costs and immobility, are not suitable in application settings. Second, most of these studies took a correlational approach, in which the focused trait was correlated with specific neural features. These correlations relied on in-sample population inference and were not necessarily generalizable to out-of-sample individuals(Dubois & Adolphs, 2016). In contrast, a predictive machine-learning inspired framework would employ cross-validation techniques to ensure out-of-sample generalizability, thus may be more desirable for application scenarios which require accurate personality predictions from novel samples.

In the present study, we propose a novel machine learning-based assessment of the Big Five personality traits using a brief electroencephalography (EEG) recording. EEG is one of the most commonly used non-invasive neuroimaging techniques and is especially suitable for application-oriented personality assessment due to its relatively inexpensive and tolerable nature(Suzuki, Hill, Ait Oumeziane, Foti, & Samuel, 2018). The premise of the proposed measure is based on a large body of previous research which shows that the Big Five traits are related to affective reactivity. For example, extroverts were shown to be more likely to experience positive emotions(Lee Anna Clark & Watson, 2008; John, Naumann, & Soto, 2008), while those scoring high on neuroticism were more inclined to experience negative emotions(Lee Anna Clark & Watson, 2008; John et al., 2008). Accordingly, studies of event-related potentials (ERPs) have shown that personality affects one’s neural response to emotional stimuli(De Pascalis, Strippoli, Riccardi, & Vergari, 2004; Y. Lou et al., 2016; Speed et al., 2015; Suzuki et al., 2018), and there are recent studies reporting distinct EEG profiles by people with high versus low level of personality traits when viewing video clips(Subramanian et al., 2018; Zhao, Ge, Shen, Wei, & Wang, 2018). However, personality inferences finer than binary levels based on brain activities have not yet been achieved. Our method aims to fill this gap by providing quantitative EEG-based predictions of the Big Five traits.

In the proposed method, participants rapidly view a series of emotional words whilst their brain activities are captured as EEG signals which are then fed to trained machine learning algorithms as features to predict their scores on each of the Big Five traits (Fig. 1A). We choose words as emotional stimuli because they are fast to process, allowing the task to be brief (~ 5 mins) and offering flexibility in application scenarios. To train the machine learning model for personality inference, and to systematically evaluate its reliability and validity, we collected data from a large sample of 196 young and healthy participants recruited from nearby universities (154 females, mean age = 21 years). Two-hundred double-character Chinese words were briefly presented in a randomized order, including 60 positive words, 60 negative words, 60 neutral words, and 20 name words. EEGs were simultaneously recorded whilst participants viewed the words. ERPs evoked in response to the three types of emotional words were extracted from the EEG recordings and used to train predictive models with a nested cross-validation approach (Fig. 1). The performances of the predictive models were evaluated using the correlations between EEG-predicted and self-reported trait scores. Furthermore, the external validity of the measure was evaluated by using the predicted traits scores to predict participants’ behavioral tendencies and life outcomes. Lastly, some of the participants completed the task again 19-78 days later, and the correlations between the predicted scores of the two time points were used to assess the test-retest reliability of the proposed EEG-based measure.

**Fig. 1.**
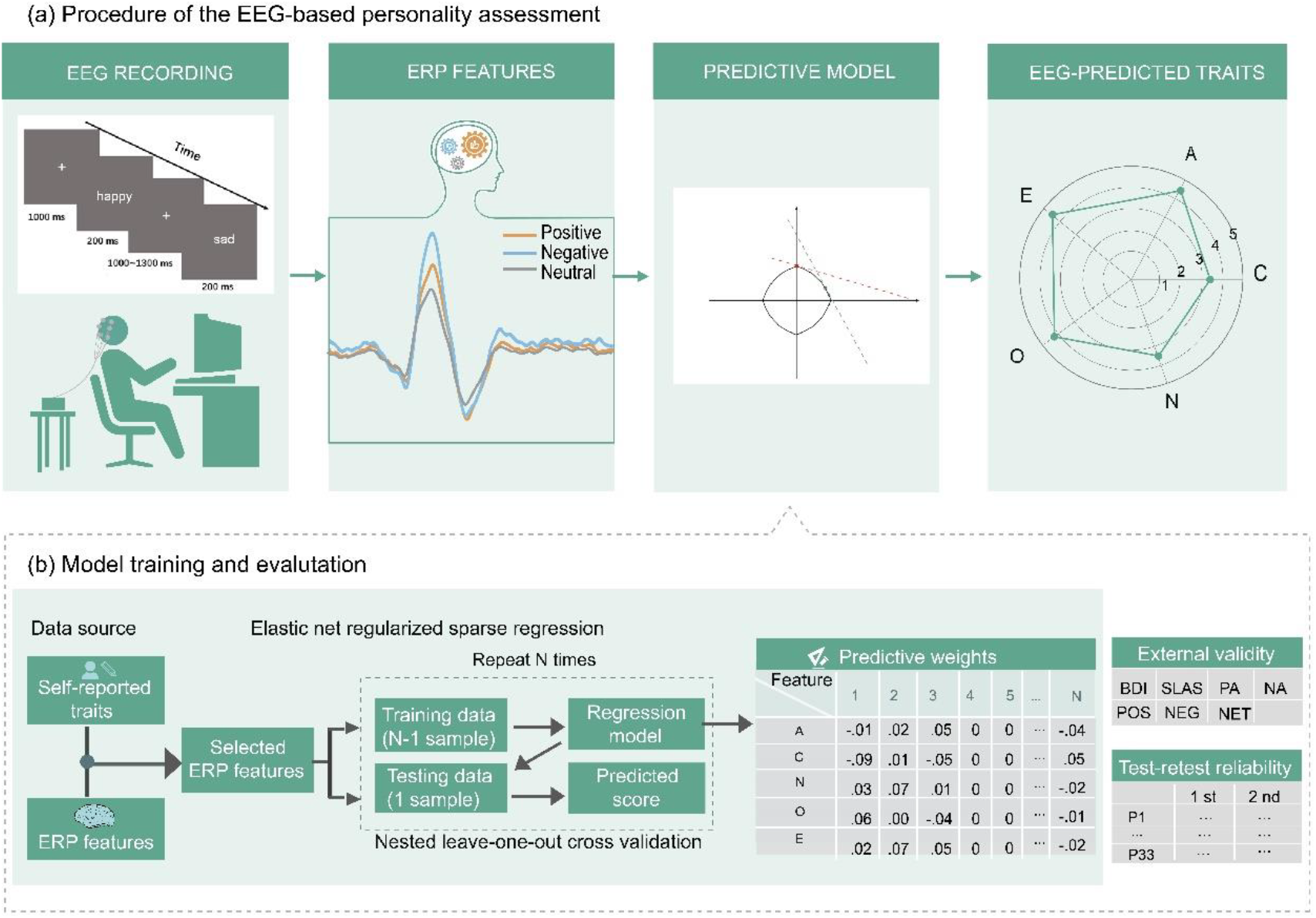
Flow chart of the data collection and overview of the model training and evaluation. (*a*) The procedure of the personality assessment task. The participants perform the word attention task while their brain activity is recorded by a portable wireless EEG system. The event-related potential (ERP) responses to positive, negative and neutral words are used as features for implementing machine learning-based predictive models. The output of the models are the predicted scores for the Big Five traits. (*b*) The procedure of model training and evaluation. Elastic net regularized sparse regression is employed, with a nested leave-one-out cross-validation strategy for feature selection and model evaluation. The models’ external validity and test-retest reliability are also assessed.

## Results

### Behavioral results

The presentation of the emotional words was randomly intermixed with 20 common Chinese name words. The participants were required to press a button when they detected a name. The mean accuracy for responding to names was 97.19 ± 5.04% and the mean response time was 522 ± 166 ms, indicating that participants were attentive during the task.

### Analyses of ERP responses

Averaged ERP responses to the word stimuli for participants with trait scores ranking in the top, middle and bottom terciles are shown in Fig. 2 for each combination of trait and word valence. The prominent ERP components elicited by the word stimuli included two positive peaks at 200-300ms and 400-500ms, and two negative peaks at 100-200ms and 300-400ms, corresponding to the emotion related ERP components of N100, P200, N400 and late positive complex (LPC)(Y. X. Lou et al., 2016; Williams et al., 2006; M. Zhang, Ge, Kang, Guo, & Peng, 2018).

**Fig. 2.**
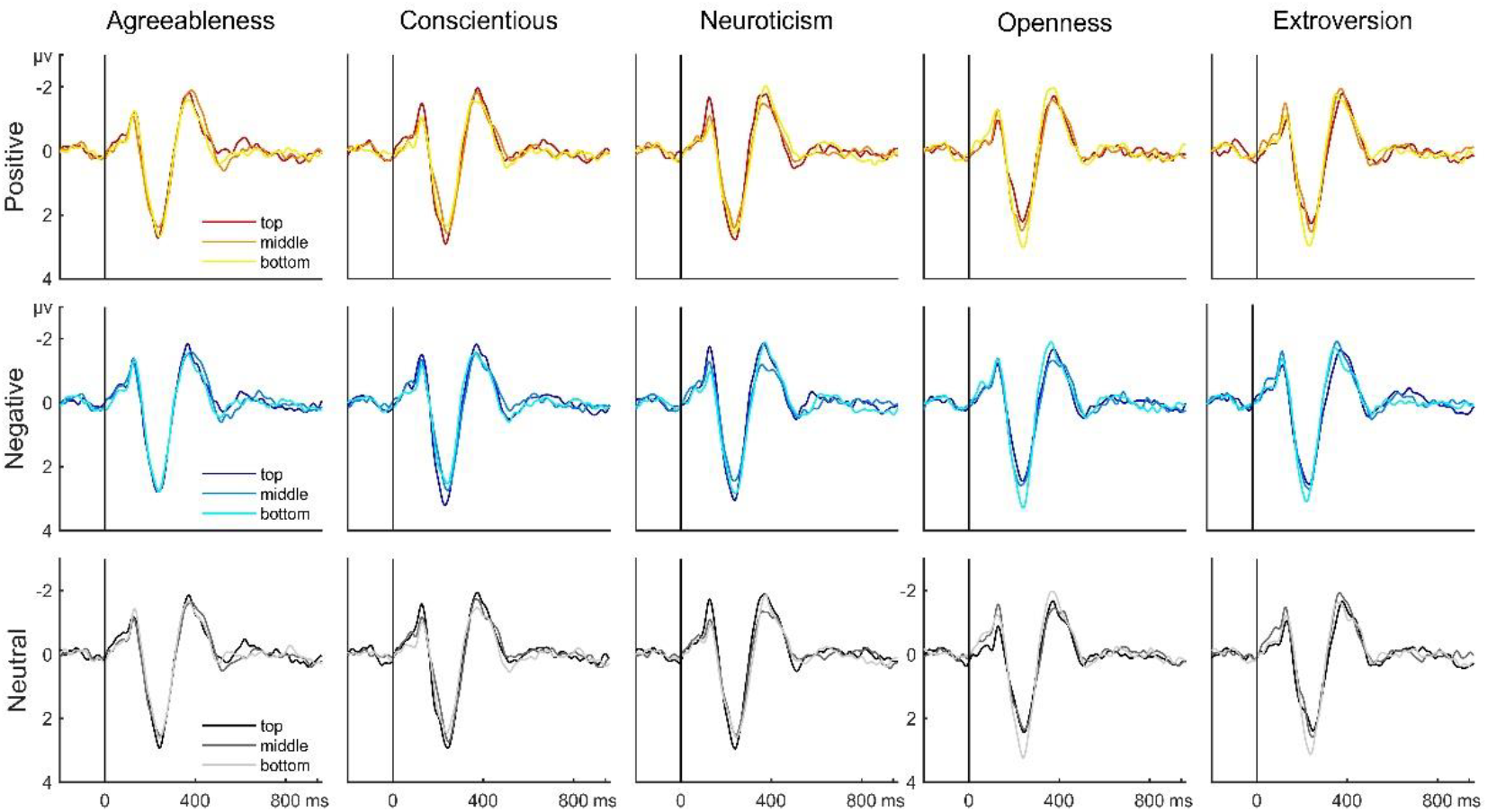
An overview of the event related potential (ERP) responses. The ERP waveforms show the average ERPs across all recording channels for the corresponding combination of trait (column) and word valence (row). The three waveforms within each subplot correspond to the ERPs averaged over the participants with the corresponding trait scores ranking in the top, middle and bottom terciles. Darker color refers to higher scores.

As a first step, we examined these components’ correlation with personality. As shown in Fig. S1, there was only one significant correlation between LPC for positive words in the temporal area and self-reported scores for agreeableness (*r* = -.18, *p* < .05). For conscientiousness, higher scores were associated with smaller LPC for neutral words in the frontal and right temporal area (*r* = -.15, -.15, respectively, both *p* < .05). For neuroticism, higher scores were associated with larger N100 for positive words in the central area (*r* = -.16, *p* < .05), larger N100 for negative words in the left temporal area (*r* = -.15, *p* < .05), larger N100 for neutral words in the frontal, central, left temporal (*r* = -.16, -.17, -.17, respectively, all *p* < .05), larger N400 for neutral words in the frontal, central, left temporal and right temporal areas (*r* = -.20, -.15, -.15, .20, respectively, all *p* < .05), larger LPC for positive words in the frontal, central, left temporal and right temporal areas (*r* = .15, .15, .17, .20, respectively, all *p* < .05). For openness, higher scores were associated with smaller P200 for positive words in the central and left temporal area (*r* = -.14, -.16, respectively, both *p* < .05). For extraversion, higher scores were associated with smaller N100 for positive words in the central area (*r* = .15, *p* < .05), smaller P200 for positive words in the central, left temporal and right temporal areas (*r* = -.16, -.19, -.16, respectively, all *p* < .05), smaller N100 for neutral words in the frontal and central areas (*r* = .18, .14, respectively, both *p* < .05), smaller P200 for neutral words in the central, left temporal and right temporal areas (*r* = -.21, -.18, -.18, respectively, all *p* < .05), and smaller N400 for negative words in the left temporal area (*r* = .15, *p* < .05).

### Predictive models of personality based on ERP responses

Participants’ ERP responses elicited by the word stimuli were used as features to train five predictive models, one for each of the Big Five traits, using a nested cross-validation approach with elastic net regularized regression analyses. To assess the predictive models’ performance, correlations were calculated between pairs of EEG-predicted and self-reported scores for each of the Big Five traits. Notably, important ERP features retained as well as finally used for the sparse-regression-based trait predictive models (see ‘Feature selection and model training’ in Methods) were located not only within the time windows of these emotion related ERP components, but also extended to the pre-stimulus periods (< 0 ms), as well as the late processing stages (> 500 ms) (Fig. 3).

**Fig. 3.**
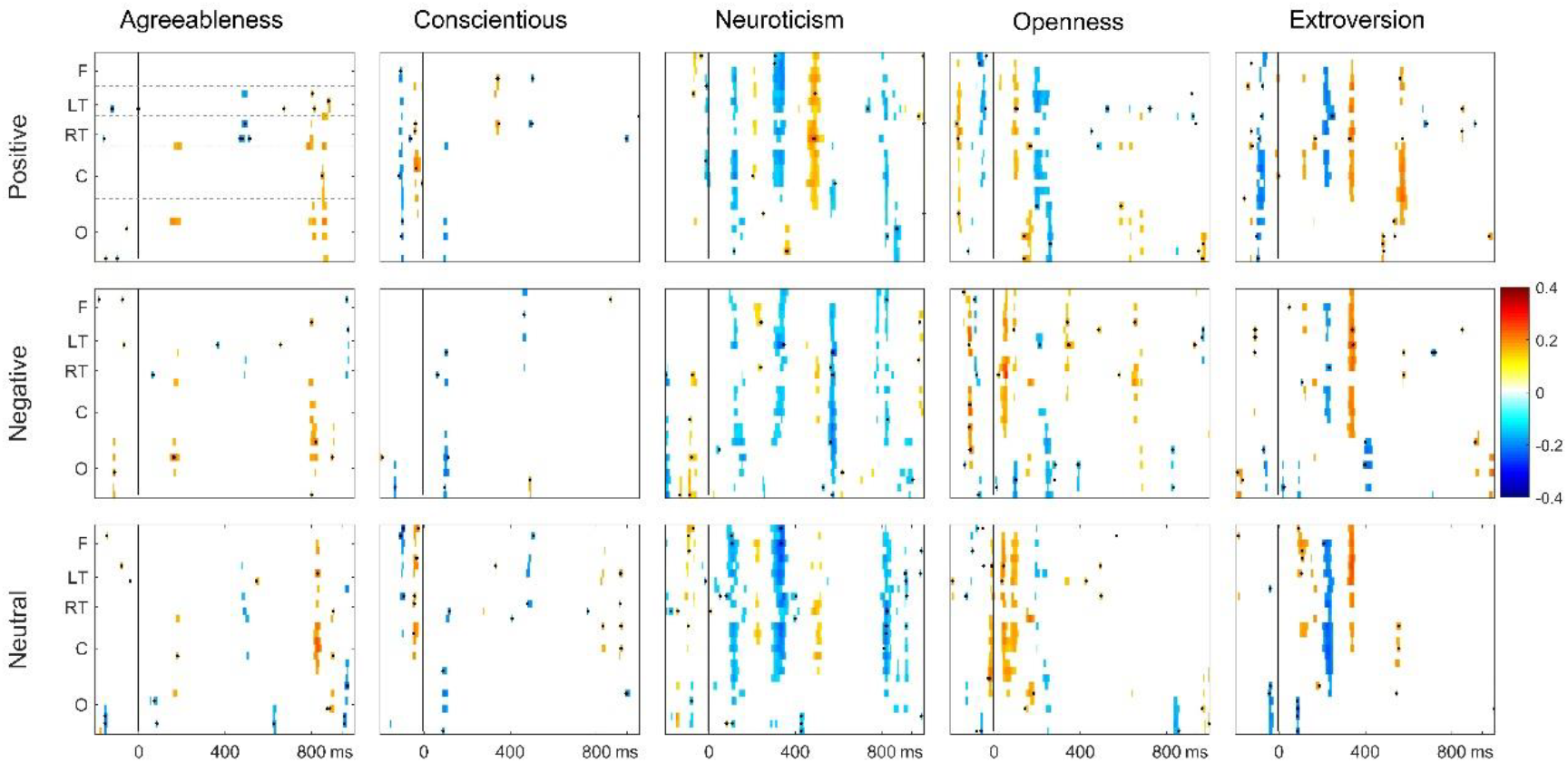
ERP features used in the trait predictive models. The colored channel by time bins demonstrate the ERP features retained for model training (*p*-value < the optimal *p*-value threshold) and the black dots mark the bins that were finally used in the elastic net regularized sparse regression model. The colors show the bivariate Pearson correlation coefficients between the ERP features at the channel-time bin and the corresponding self-report trait scores. EEG channels array are Fp1/2, Fz, F3/4, F7, FC5, T3, CP5, F8, FC6, T4, CP6, FC1/2, Cz, C3/4, CP1/2, P3/4, Pz, PO3/4, Oz, O1/2, organized in five ROIs: frontal area (F), left temporal area (LT), right temporal area (RT), central area (C), occipital area (O). See ‘Feature selection and model training’ in Methods for details.

The predictive models achieved significant correlations between the predicted and self-reported trait scores (Fig. 4). Specifically, Pearson correlations for agreeableness, conscientiousness, neuroticism, openness and extroversion were .47, .61, .49, .48, and .53, respectively (all *p* < .001, *N* = 196).

**Fig. 4.**
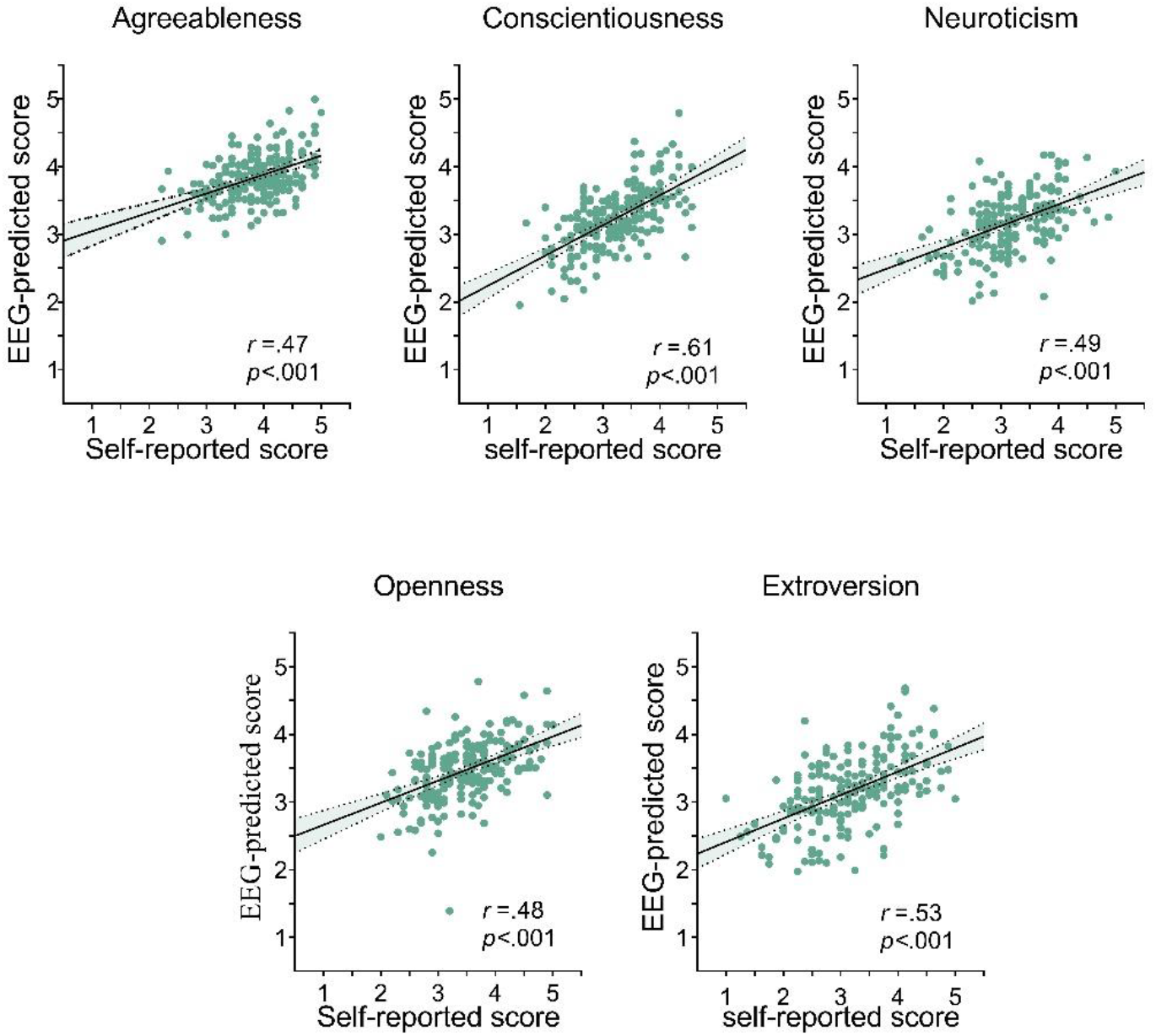
Scatterplots for the correlations between the predicted and self-reported trait scores. Each dot represents the scores from one participant (for each plot, *N* = 196). The predicted score for each dot was obtained by using a nested cross-validation approach with the predictive model trained with the remaining samples excluding the to-be-predicted sample.

For 127 of the 196 participants, the mean participant-wise absolute difference between the predicted and self-reported scores (averaged over the absolute differences from the five traits) were less than 0.5 on a 5-point scale (Fig. 5a, mean differences across participants = 0.45±0.1 8). In addition, the histogram of the correlation coefficients between the 5-dimensional EEG-predicted personality trait constructs and the self-reported counterpart for each individual participant shows a clear tendency towards high correlation values (Fig. 5b): 139 out of the 196 participants showed correlations higher than .5 (average correlation *r* = .59 ± .37). The high correlation values indicate that these five predictive models together can reliably reflect the relatively high and low of the participants’ personality scores.

**Fig. 5.**
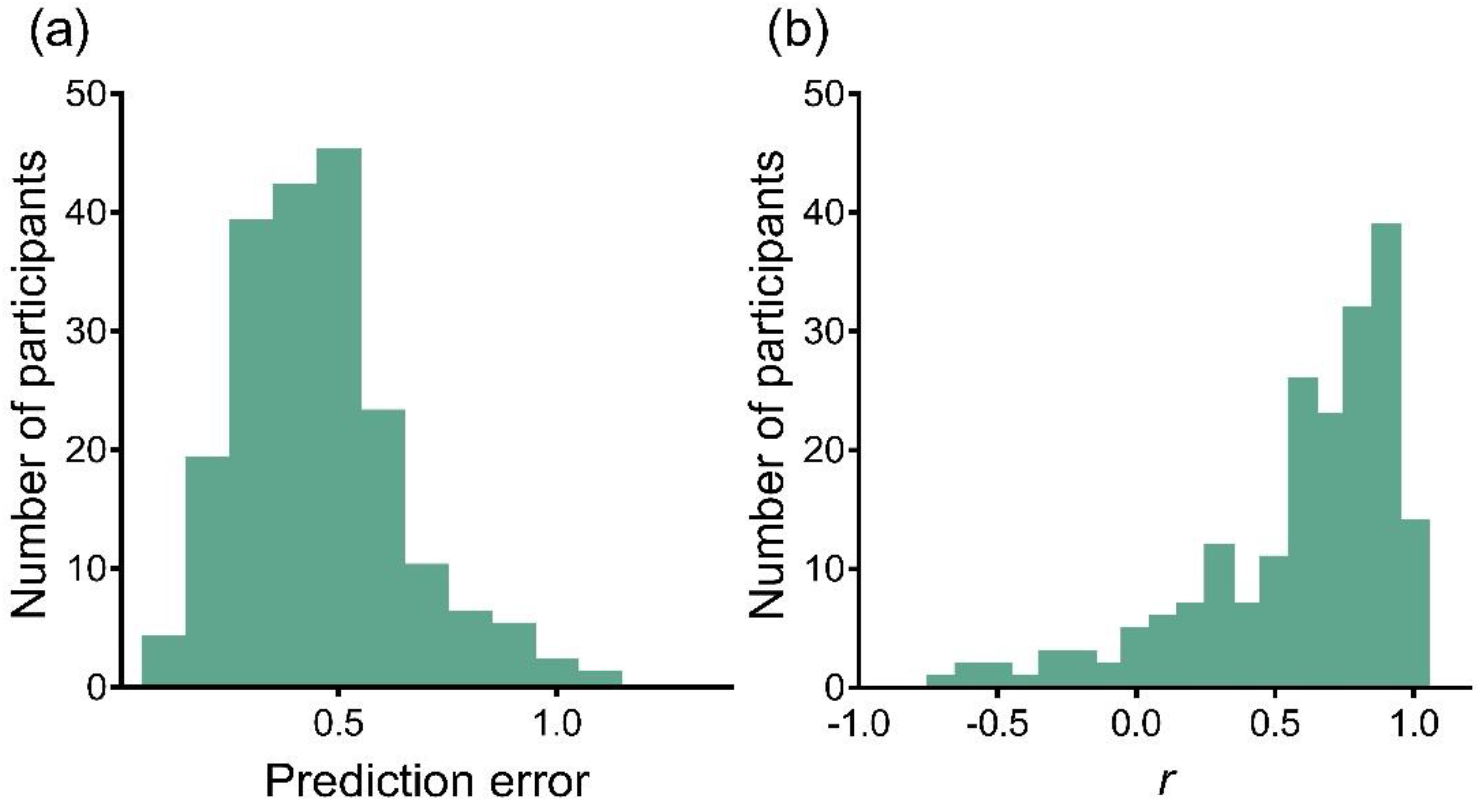
Evaluation of the predicted scores. (*a*) A histogram of the participant-wise prediction errors (i.e. the mean absolute difference between the EEG-predicted scores and self-reported scores). (*b*) A histogram of the participant-wise correlations of the 5-dimension personality constructs between the EEG-predicted scores and self-reported scores.

### External validity

After the task, a subsample of the participants also completed one or two sets of measures for assessment of external validity. First, a subsample of the participants completed questionnaires for indices of psychological adjustment, including life satisfaction (SLAS, *N* = 135), positive affects (PA, *N* = 111), negative affects (NA, *N* = 111), and symptoms of depression (BDI, *N* = 111), which have been shown to be predicted by personality scores in previous studies(Cloninger, Svrakic, & Przybeck, 2006; González Gutiérrez, Jiménez, Hernández, & Puente, 2005; Larsen & Ketelaar, 1991; Strickhouser, Zell, & Krizan, 2017). Second, 60 participants also watched a series of emotional video clips and rated the valence of each clip. The averaged valence ratings for the positive (POS), negative (NEG), and neutral (NET) clips were used as measures of their affective responses to emotional stimuli. For each of the seven indices, two separate regression models were built using the EEG-predicted and self-reported trait scores, and external validity was assessed using the regression model fitting *R* values. For the four indices of psychological adjustment as well as the valence rating for positive video clips, the self-reported trait scores achieved higher predictive power than the EEG-predicted trait scores. However, for the experienced emotional valences the neutral and negative video clips, the EEG-predicted scores were able to achieve slightly higher predictive powers than self-reported scores (Fig. 6).

**Fig. 6.**
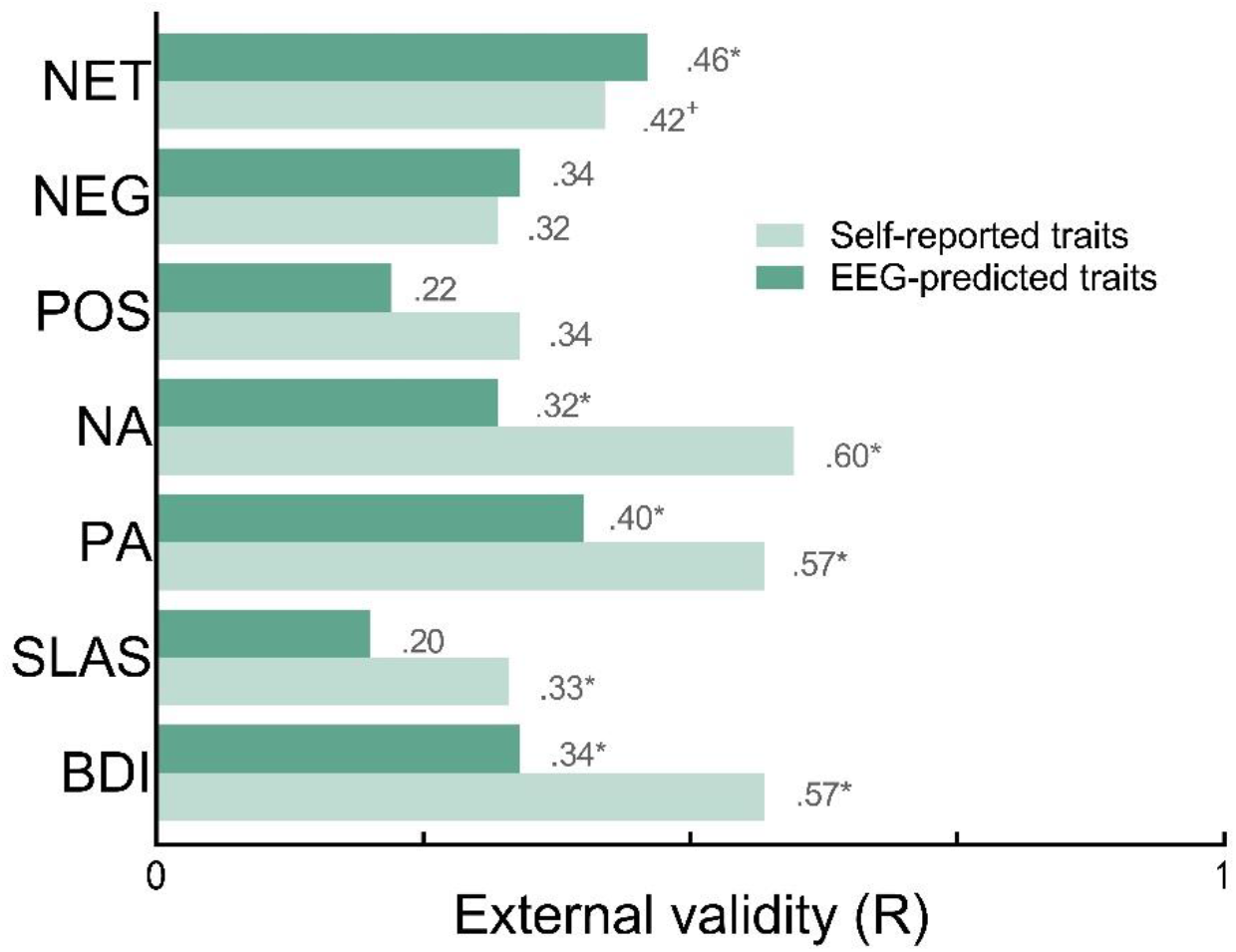
External validity of the EEG-predicted and self-reported trait scores. The dark green and light green bars show the predictive powers of EEG-predicted and self-reported trait scores for a certain behavior or life outcome index as reflected in regression model fitting r values. NET, NEG, POS are participants’ ratings of the valence of neutral, negative and positive video clips, NA and PA are self-reported scores of negative and positive affects; SLAS is the self-reported score of Satisfaction with Life Scale; BDI is the self-reported score of Beck Depression Inventory. See Table S2 for detailed results.

### Test-retest reliability

Temporal correlations were calculated for each of the predicted and self-reported trait scores from the subsample of the participants (*N* = 33) who completed the task for a second time 19-78 days later. The self-reported trait scores showed adequate to good test-retest reliability (*r* = .86, .67, .65, .76 and .79 for agreeableness, conscientiousness, neuroticism, openness and extroversion, respectively). The predicted scores’ test-retest reliability, except for neuroticism, were lower than the self-reported scores (*r* = .51, .31, .67, .50 and .58 for agreeableness, conscientiousness, neuroticism, openness, and extroversion, respectively). A closer look at the data suggested that the extremely low reliability of conscientiousness was largely due to two outliers. After these two were excluded, the reliability increased to .65. Participant-wise analyses revealed that the average of the mean score difference over the five traits was 0.27 ± 0.15, and the average 5-dimension construct-based correlation was .67 ± .31 (Fig. 7).

**Fig. 7.**
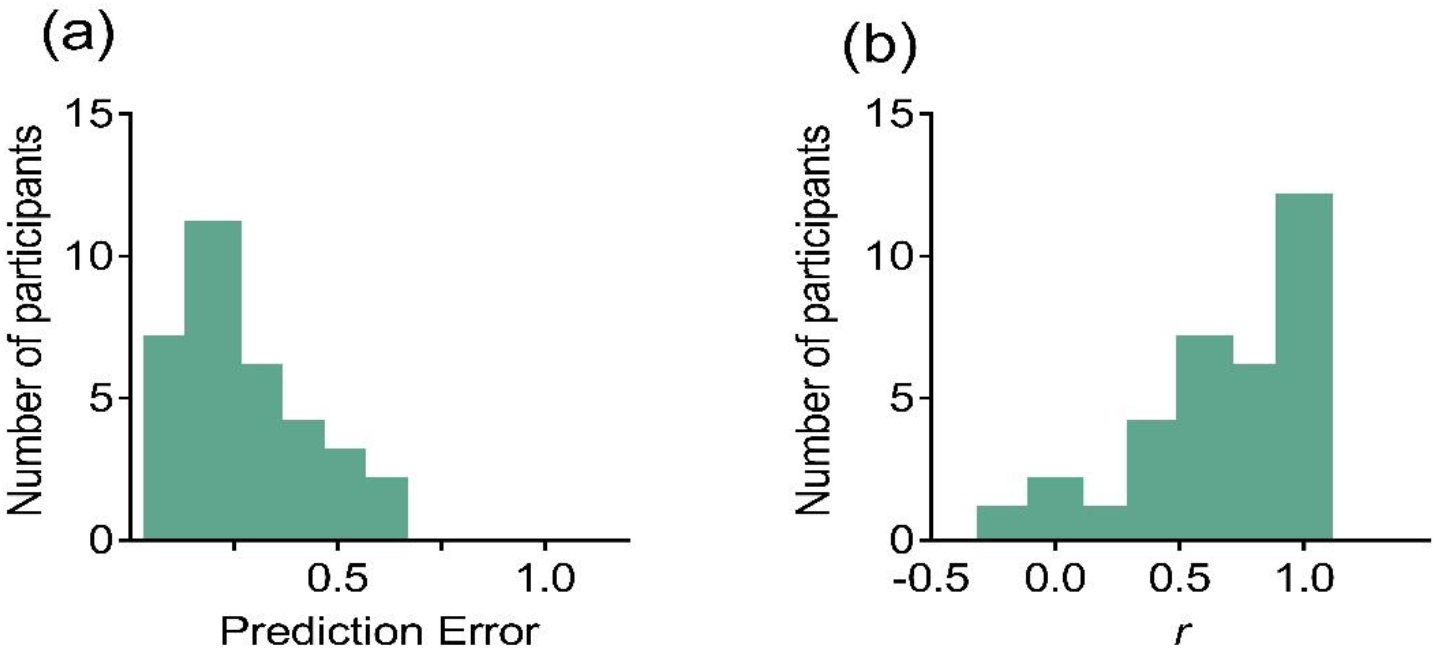
Evaluation of the test-retest reliability. (*a*) A histogram of the participant-wise test-retest errors across two data collections. (*b*) A histogram of the participant-wise correlations of the scores of the 5-dimension personality constructs between the two data collections.

## Discussion

Our results for the first time demonstrate the feasibility of combining machine learning and EEG recordings to make indirect yet fairly accurate quantitative predictions about an individual’s personality. The correlations between the predicted and self-reported scores (.47-.61) were comparable to previous studies using digital footprints as input features(Wu et al., 2015). Furthermore, the EEG-predicted scores could significantly predict several indices of psychological adjustment, even though their predictive powers were lower than those of the self-reported scores. The better performances of the self-reported trait scores might be partially attributed to the fact that psychological adjustment was also measured with self-reported scales, and common-method bias may have inflated the correlations among them(Podsakoff, Mackenzie, Jeong-Yeon, & Podsakoff, 2003). For outcomes like affective responses to video clips, the EEG-predicted trait scores achieved slightly better predictive powers than the self-reported scores, demonstrating their usefulness in predicting real-world affective experiences. While producing results comparable to self-reported measures, the proposed method does not require the participant to report his/her own personality explicitly, thus is less susceptible to faking. Also, the task is brief in time and has been tested with a portable EEG system, making it useful for application-oriented personality assessment.

Even though we primarily focused on developing a new method for personality assessment, a closer look at the correlation between personality and the temporal and spatial patterns of standard ERP features may also shed some light into the question of the neurophysiological basis of personality. Firstly, in general, extroversion and neuroticism were associated with more ERP components, which is consistent with the previous finding that these two traits more closely connect to emotions(L. A. Clark, 2005; L. A. Clark, Watson, & Mineka, 1994; Watson, Clark, & Harkness, 1994). Secondly, there were significant correlations between ERP responses for positive words in the temporal area and self-reported scores for agreeableness and openness. These results are consistent with previous studies reporting that these two traits are associated with positive affects(Holtgraves, 2011; Letzring & Adamcik, 2015; Ready & Robinson, 2008), and that agreeableness is closely associated with the temporal regions responsible for social information processing(DeYoung et al., 2010; B. W. Haas et al., 2015; Haas, Ishak, Denison, Anderson, & Filkowski, 2015). Finally, for conscientiousness, we observed a diminished LPC for neutral words for the participants with higher conscientiousness scores, which may support the hypothesis that conscientiousness reflects a tendency to inhibit impulses and feel calmness(Fleming, Heintzelman, & Bartholow, 2016; John et al., 2008). Nonetheless, these correlations were generally weak in magnitude (.15-.21), making it difficult to make accurate individualized inferences. The machine learning approach, on the other hand, simultaneously took multiple neural features into considerations and produced more reliable individualized predictions. Furthermore, the cross-validation techniques used in the development of the predictive algorithm ensures greater out-of-sample generalizability(Dubois & Adolphs, 2016), thus could be more useful for application purposes such as personnel selection.

It might also be worthwhile to examine the predictive performances of models using ERP responses from only a single condition (positive, negative or neutral words). In general, these models’ performances were sub-par compared to models using data from all three conditions (Fig. S2). With single condition models, the best performing condition for extroversion was the positive condition. This is consistent with previous studies which have found that extroverts are more closely associated with positive emotions(Canli et al., 2001; Lucas, Le, & Dyrenforth, 2008; Srivastava, Angelo, & Vallereux, 2008; L. Wang, Shi, & Li, 2009; Yuan, He, Lei, Yang, & Li, 2009; Yuan et al., 2012). For openness and neuroticism, the models in three conditions had similar performance. This is also consistent with previous studies which have suggest that both dimensions are associated with the processing of stimuli of various valences(John et al., 2008), (Bartussek, Becker, Diedrich, Naumann, & Maier, 1996; Gray, 1981). In the models for conscientiousness and agreeableness, there was better performance in the neutral condition. These results are consistent with the definition of the two dimensions, which are less related to emotional reactivity(John et al., 2008). Even though we designed the measure based on the Big Five’s relationship with the processing of emotional stimuli, the predictive weights of the neutral features suggest that non-affective processes may also contribute to the predictive models’ performances.

Interestingly, when taking a closer look at the temporal aspects of feature selection, there were selected features from the pre-stimulus period for all the predictive models. The nature of pre-stimulus ERP components has long been a topic of discussion. While the ERP signals recorded before the onset of stimuli have traditionally been considered as “baseline” and not included in data analysis, there is emerging evidence to suggest that there are functional implications for pre-stimulus activity(Falkenstein, Hoormann, Christ, & Hohnsbein, 2000; Lazzaro, Gordon, Whitmont, Meares, & Clarke, 2001). The inter-trial variability of the pre-stimulus activity has been repeatedly been reported as being related to one’s cognitive states(Bode et al., 2012; Ikumi, Torralba, Ruzzoli, & Soto-Faraco, 2019; Lou, Li, Philiastides, & Sajda, 2014; Polich & Kok, 1995). As the mean amplitude of the pre-stimulus period was subtracted before the analysis, our results suggest a possible contribution from the fluctuation of the baseline activity rather than its absolute amplitude. In addition, our study found associations between the inter-participant variability of the baseline ERP responses and one’s trait scores. Therefore, our findings extend existing findings by suggesting that baseline activity might provide information about one’s dispositional tendencies. However, it should be noted that the above discussions based on feature selection are mostly speculative. More theoretical and empirical works are needed to clarify the psychological and neural mechanism.

The test-retest reliabilities for agreeableness, openness, and extroversion of the proposed EEG measure were in the range of. 5-.7. While these results were generally lower than the self-reported counterpart (in the range of. 7-.8), our findings are comparable, if not better, than the existing studies on the stability of ERP responses(Ip et al., 2018; Segalowitz & Barnes, 1993). According to previous studies, the reliability of EEG and ERP was affected by various variables, such as age of participants(Alperin, Mott, Rentz, Holcomb, & Daffner, 2014), recording intervals(Sandman & Patterson, 2000), state and other factors(Ip et al., 2018; Segalowitz & Barnes, 1993). In our study, one possible source of error may have been if the EEG cap aligned slightly differently between the two data collection sessions. Thus, the positions of the electrodes may have deviated slightly, introducing additional noise into the predictive models. In addition, a systematic evaluation and control of the participant’s general cognitive state should have been conducted, as it could substantially affect the emotional ERP responses(Jiang et al., 2017). Further studies are necessary to elucidate these issues, especially focusing on the participants with low test-retest reliabilities.

As a final, but note-worthy comment, while the present study was conducted using a wet electrode based EEG system, recent advances in EEG recording techniques on electrode materials and designs, hardware improvements and system optimization have shown the potential to greatly improve the usability of EEG devices to a general user population(Lühmann, Wabnitz, Sander, & Müller, 2017; Siddharth, Patel, Jung, & Sejnowski, 2018; F. Wang, Li, Chen, Duan, & Zhang, 2016). The proposed EEG based personality measure is expected to be readily applicable in many practical scenarios, serving as a promising alternative to conventional personality questionnaires in the near future.

## Materials and Methods

### Participants

One hundred and ninety-six young participants (154 females, mean age = 21 years, range 18-28 years) from Tsinghua University and China Women’s University took part in the study. All of them had normal or corrected-to-normal vision. Informed consent was obtained from all participants. The study was conducted in accordance with the Declaration of Helsinki and approved by the local Ethics Committee of Tsinghua University.

### Materials

One hundred and eighty double-character Chinese words were employed as the stimuli, including 60 positive-emotion words, 60 negative-emotion words, and 60 neutral-emotion words (see Table S2 for the full list). All words were selected from the Chinese Affective Words System(Y. N. Wang, Zhou, & Luo, 2008; Q. Zhang, Li, Gold, & Jiang, 2010). According to their valence, we choose the top 20 most pleasant adjectives, nouns and verbs as positive-emotion words (mean valence rating 7.43±0.16 on a 9 -point Likert scale), the top 20 least pleasant adjectives, nouns and verbs as negative-emotion words (mean valence 2.38±0.21), the median 20 pleasant adjectives, nouns and verbs as neutral words (mean valence 5.52±0 .71). In addition, 20 double-character common Chinese names were selected as non-emotional stimuli for the behavioral task.

The Chinese version of the Big Five Inventory (BFI)(Carciofo, Yang, Song, Du, & Zhang, 2016) was used to measure participants’ personalities. The questionnaire is a 5-point Likert scale including 44 items, 8 measures of extraversion, 9 measures of agreeableness, 9 of measures conscientiousness, 8 measures of neuroticism and 10 measures of openness. The internal consistency coefficients were good for every dimension in the current study (alpha: extraversion = .89, openness = .85, neuroticism = .84, conscientiousness = .82, agreeableness = .79).

### Experimental procedure

The experiment was carried out in a regular laboratory environment without any electrical shielding. There was ambient illumination from ceiling lights. The stimuli were displayed on a 22-inch LCD monitor (DELL, USA) with a 60 Hz refresh-rate. The participants sat in a comfortable chair approximately 60 cm away from the monitor screen.

The participants first filled in the BFI questionnaire prior to the start of the experiment. The main experiment consisted of 200 trials (Fig. 1). Within each trial, one double-character Chinese word was presented for 200 ms, followed by an intertrial interval of a random length in the range 1000-1300 ms. All words were presented in white against a black background. Words were presented in the center of the computer screen, with a size of 1.5° by 2.0° (horizontal by vertical, measured in visual angle) per character and a 0.75° center-to-center distance between the characters. The order of the presentation was randomized for each participant. The participants were asked to focus on the words and press the Down Arrow key on the computer keyboard when they detected a Chinese name. The duration of the EEG recording was about 5 minutes per participant (excluding the EEG preparation time). Presentation of the stimuli and collection of the behavioral responses were programmed in MATLAB (The Mathworks, USA) using the Psychophysics Toolbox 3.0 extensions(Brainard, 1997; Kleiner et al., 2007; Pelli, 1997).

### EEG recordings

A portable wireless EEG amplifier (NeuSen.W32, Neuracle, China) was used for data recording at a sampling rate of 250 Hz. EEG data were recorded from 28 electrodes positioned according to the international 10-20 system (Fp1/2, Fz, F3/4, F7/8, FC1/2, FC5/6, Cz, C3/4, T3/4, CP1/2, CP5/6, Pz, P3/4, PO3/4, Oz, O1/2) and referenced to linked mastoids with a forehead ground at AFz. Electrode impedances were kept below 10 kOhm for all electrodes throughout the experiment.

### EEG preprocessing

All EEG data analyses were performed using MATLAB with the Fieldtrip toolbox(Oostenveld, Fries, Maris, & Schoffelen, 2011). The continuous EEG data were first band-pass filtered at 1-30 Hz. Artifacts due to eye movement, muscle movement, and other possible environmental noises were removed using independent component analysis (ICA). On average, 1-3 artifact related independent components (ICs) per participant were manually identified and excluded. The remaining ICs were then back-projected onto the scalp EEG channels to reconstruct the cleaned EEG data. EEG data were then segmented into 1.2-sec trials from 200 ms pre-stimulus to 1000 ms post-stimulus. Trials with non-emotional stimuli (i.e., Chinese names) were excluded from further analysis. Trials with peak-to-peak voltage changes exceeding ±150 mV in any recording electrode were also rejected to avoid possible artifact contamination. On average, the number of rejected trials per participant was less than 10. The artifact-free trials were then averaged for each emotional category (i.e., positive, negative and neutral) and baseline corrected using the average of the 200 ms pre-stimulus data, resulting in three ERP waveforms per participant.

### ERP component analysis

This research analyzed the potentials of the N100, P200, N400 and LPC components across different sets of electrodes. The mean amplitude of all ERPs component was calculated in five ROIs and four time windows (frontal area: Fp1/2, Fz, F3/4; central area: FC1/2, Cz, C3/4, CP1/2; left temporal area: F7, FC5, T3, CP5; right temporal area: F8, FC6, T4, CP6; occipital area: P3/4, Pz, PO3/4, Oz, O1/2; time windows: 100–140 ms for N100; 200–280 ms for P200; 320-400 ms for N400; and 460-540 ms for LPC).

Pearson’s correlations were computed between the mean amplitudes of N100, P200, N400 and LPC components for different emotional words (positive, negative, neutral) and self-reported scores, with uncorrected *p-*values reported.

### Feature selection and model training

The processed data were used as features for building regression models for the prediction of the five trait scores. The averaged multichannel ERP responses to positive, negative and neutral words yielded 3 (emotion: positive, negative and neutral) × 28 (EEG channels) × 300 (sample points comprising 1.2 s at a sampling rate of 250 Hz) = 25, 200 features per sample (participant). As the feature dimensions were much larger than the sample size (i.e., 196 participants), it was necessary to perform feature selection for enhancing the stability and generalizability of the regression models(Bermingham et al., 2015). Following previous neuroimaging studies(Cui, Xia, Su, Shu, & Gong, 2016; R. T. Jiang et al., 2018; Rosenberg, Hsu, Scheinost, Todd Constable, & Chun, 2018), we applied a nested leave-one-out cross-validation (nested-LOOCV) strategy, including an outer and an inner loop. The procedure was performed separately for each of the five traits.

The outer loop performed the overall evaluation of the models generated by the inner loop. By leaving out one sample (participant) at a time, the remaining 195 samples were used as the training set to build 196 regression models (with the self-reported scores of one trait as the dependent variable). These regression models were then applied to the left-out sample to obtain 196 predicted personality scores. The Pearson’s correlation coefficient between these predicted scores and their corresponding self-reported scores was used to quantify the effectiveness of the models. The model with the highest correlation coefficient was considered the best-performing model for further analyses.

The inner loop focused directly on feature selection. Here all analyses were performed using 195 samples from the training set as described in the outer loop procedure. The features were initially selected by thresholding the features according to the *p*-values of their bivariate Pearson correlations with the self-reported personality scores (performed separately for each personality score). By varying the *p*-value threshold from. 01 to. 15 with a step of .01, different numbers of features were retained and used for a series of regression analyses. Considering the possible occurrence of a high feature dimension problem in these conditions, a sparse regression analysis method was employed, using elastic net regularization with the alpha parameter set to 0.75(Zou & Hastie, 2005). All models were first evaluated using the outer loop, and the optimal *p*-value was subsequently decided. The changes of cross-validated correlation coefficients as a function of the *p*-value thresholds is shown in Figure S3. The optimal *p*-values for the five personality models were .03, .02, .08, .05 and .02 for Agreeableness, Conscientiousness, Neuroticism, Openness and Extroversion respectively. Correspondingly, 74, 56, 90, 90, and 70 features on average were retained for the 196 predictive models of the five traits, respectively.

The procedure is also briefly illustrated in Fig. 1 (lower panel). The LASSO method was implemented using the Statistics and Machine Learning Toolbox provided by MATLAB (The MathWorks, USA).

### Evaluation of the predicted scores

Firstly, the model performance was assessed by correlating the predicted trait scores with the self-reported scores (Fig. 4), computing prediction errors (the mean absolute difference between the predicted and self-reported scores for each trait, Fig. 5a) and computing participant-wise correlations (the correlations of the 5-dimension personality constructs between the EEG-predicted scores and self-reported scores, Fig. 5b).

Secondly, the external validity of the measure was assessed by comparing the predictive power of the predicted scores to the self-reported scores (Fig. 6). A subsample of participants completed a number of self-reported measures of indices of psychological adjustment, including the Satisfaction with Life Scale(Xiong & Xu, 2009) (N = 135), Beck Depression Inventory(Shek, 1990) (N = 111), and Positive and Negative Affects Scale(Huang, Yang, & Li, 2003) (N = 111). Another sixty participants watched 28 emotional videos including 12 positive clips (i.e., amusement, joy, inspiration, and tenderness), 12 negative clips (i.e., anger, disgust, fear, and sadness) and 4 neutral clips, all of which were selected based on standardized emotion ratings from three established emotional video datasets(Hu et al., 2017; Liu et al., 2018; Schaefer, Nils, Sanchez, & Philippot, 2010). After watching each of the clips, participants reported their experienced emotional valence of the video. The average valence of all positive (negative/neutral) clips was calculated as the final indices of positive (negative/neutral) experiences. The information of the video clips is provided in Table S3.

Finally, to assess the test-retest reliability of the models, 33 participants participated in the experiment twice, with a time interval of from two weeks to two months (mean interval 41 days, range 19-78 days). Correlations were computed between the predicted scores from the two data collection sessions. The test-retest reliability of self-reported scores was calculated in the same way. Meanwhile, prediction errors (the mean absolute differences between the predicted scores from the two data collection sessions, Fig. 7a) and participant-wise correlations (the correlations between the predicted scores from the two data collection sessions, Fig. 7b) were also computed.

## Acknowledgements

This work is supported by National Key Research and Development Plan (2016YFB1001200), National Science Foundation of China (U1736220), MOE (Ministry of Education China) Project of Humanities and Social Sciences (17YJA190017), National Social Science Foundation of China (17ZDA323), and Tsinghua University School of Social Sciences & Institute for Data Science.

We acknowledge Zhonghui Wang, Fei Dong, Xinyue Bi and Meimei Liu for help in data collection.

## Author information

### Contributions

D.Z. and F.W. developed the study concept and design. W.L. and C.W. developed the study stimuli. W.L., C.W., X.H., and J.C. collected the data. W.L. analyses and interpreted the data under the supervision of D.Z.. W.L. drafted the manuscript. S.F., X.H., and J.C. discussed the results and commented on the manuscript. D.Z. and F.W. provided critical revisions.

### Competing interests

The authors declare no competing interests.

