## Supplemental Information for "Quantitative Personality Predictions from a Brief EEG Recording"

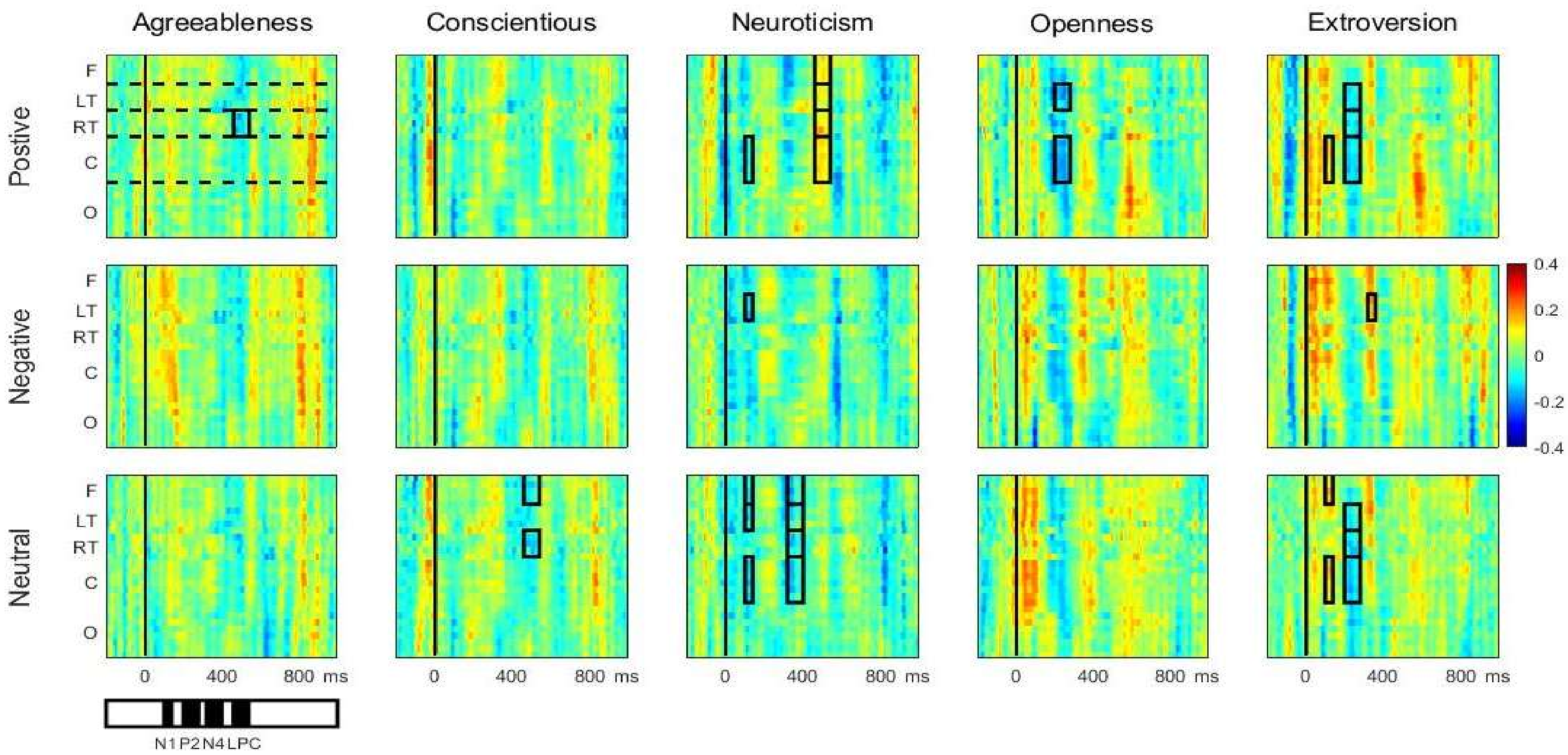


**Fig. S1.** Correlations between ERP response and self-reported personality scores. The channel by time images indicate the correlation between each ERP feature and self-reported trait scores. The rectangle area in these charts show the significantly correlation between ERP components (N1, P2, N4, LPC) and self-reported trait scores. F, LT, RT, C, O respectively represent frontal area, left temporal area, right temporal area, central area and occipital area (see 'ERP component analysis' in Method for details).

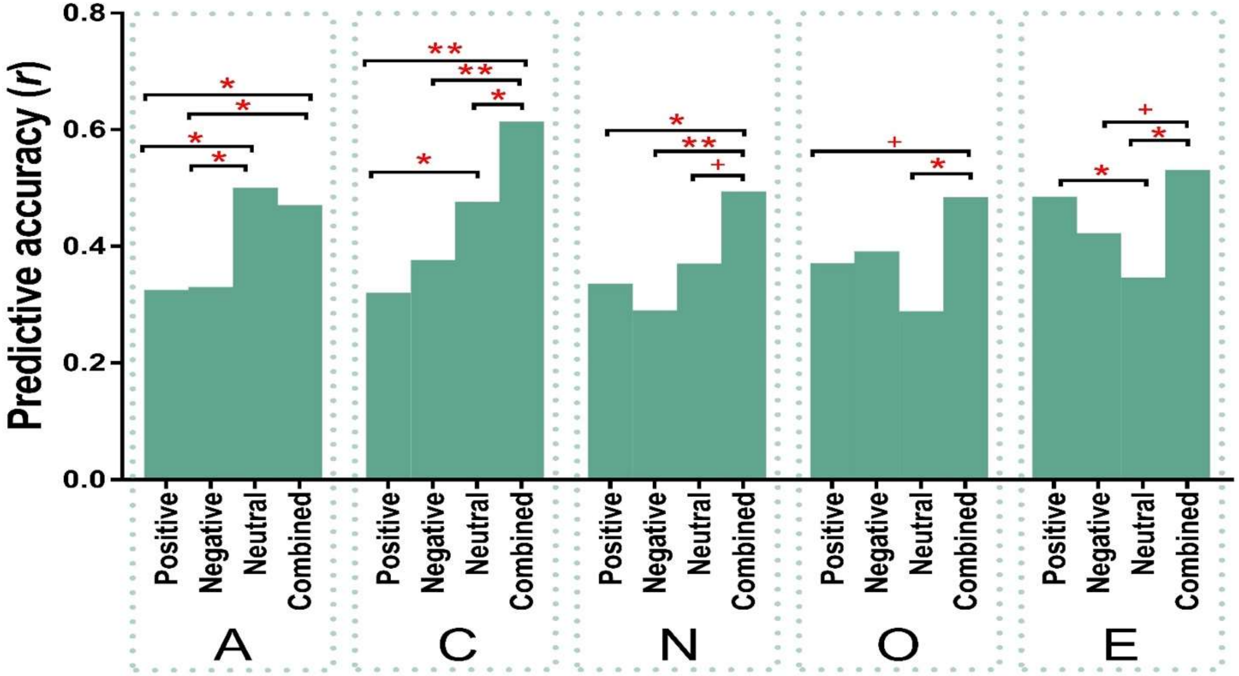


**Fig. S2.** The predictive performance using different feature sets. Each bar shows the performance of one predictive model. Positive, Negative, and Neutral refer the predictive models using features from ERPs to positive, negative and neutral words only. Combined refers to our main results with all the ERP features used. A, C, N, O, and E respectively indicate agreeableness, conscientiousness, neuroticism, openness and extroversion.

+: $p_{one\text{-}tailed} < .1$, *: $p_{one\text{-}tailed} < .05$, **: $p_{one\text{-}tailed} < .01$.

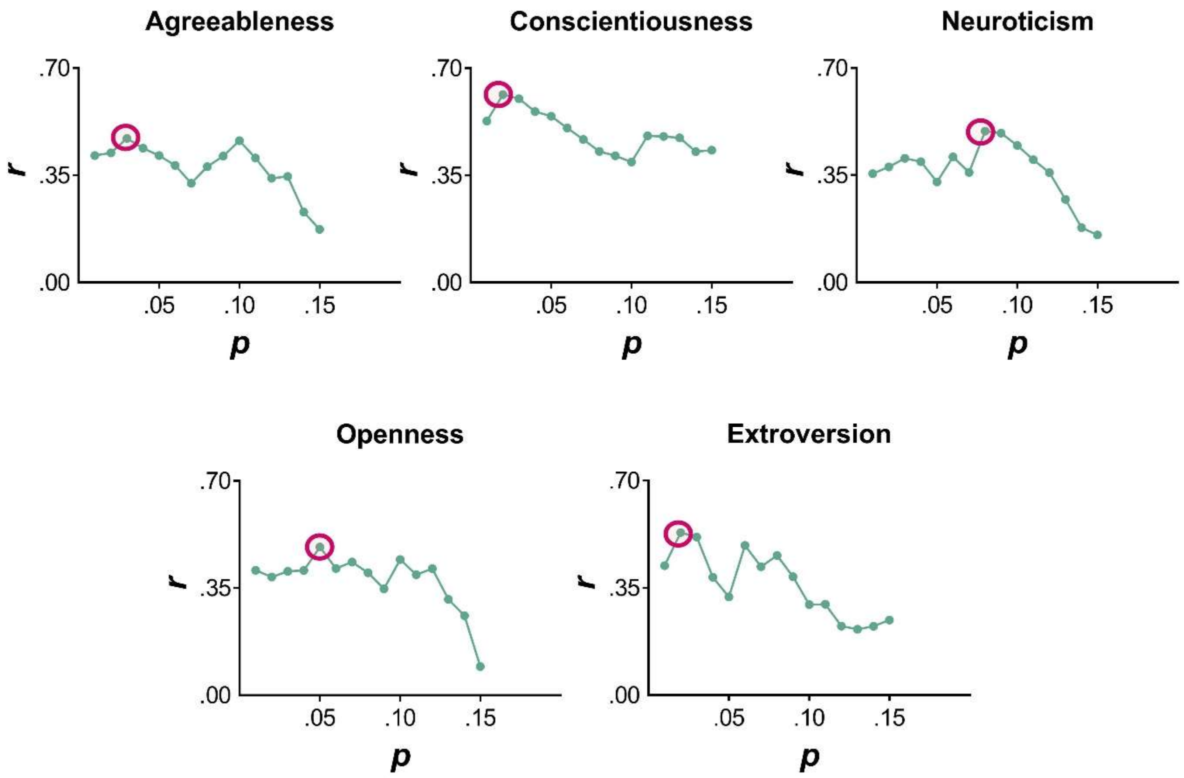


**Fig. S3.** The changes of cross-validated correlation coefficients as a function of the *p*-value thresholds. The *p*-value thresholds with best performances (marked in red circles) were chosen for implementing the regression models.

**Table S1.** The words list used in the present study. English translation is provided in brackets.

| Positive words ($N = 60$) | Neutral words ($N = 60$) | Negative words ($N = 60$) | Common Chinese Names ($N = 20$) |
|---|---|---|---|
| 深情（affectionateness） | 成年（adult） | 死刑 (death penalty) | 张伟 (Wei Zhang) |
| 真心 (sincerity) | 地位 (status) | 地狱 (hell) | 王芳 (Fang Wang) |
| 宝石 (jewel) | 首相 (premier) | 恶魔 (demon) | 李娜 (Na Li) |
| 母亲 (mother) | 银行 (bank) | 叛徒 (traitor) | 张敏 (Min Zhang) |
| 朋友 (friend) | 规律 (rule ) | 凶手 (murderer) | 王静 (Jing Wang) |
| 信心 (confidence) | 睡眠 (sleep) | 罪犯 (criminal) | 刘伟 (Wei Liu) |
| 才华 (talent) | 皇帝 (emperor) | 魔鬼 (devil) | 张丽 (Li Zhang) |
| 友谊 (friendship) | 人情 (relationship) | 罪恶 (evil) | 李强 (Qiang Li) |
| 鲜花 (flower) | 军人 (soldier) | 恶意 (venom) | 王敏 (Min Wang) |
| 美德 (virtue) | 体操 (gymnastics) | 小偷 (thief) | 王磊 (Lei Wang) |
| 情侣 (couple) | 权力 (power) | 耻辱 (humiliation) | 刘洋 (Yang Liu) |
| 成就 (achievement) | 人民 (people) | 灾害 (calamity) | 王勇 (Yong Wang) |
| 天堂 (heaven) | 眼睛 (eye) | 火灾 (conflagration) | 李军 (Jun Li) |
| 美女 (beauty) | 都市 (city) | 奴才 (slave) | 张勇 (Yong Zhang) |
| 英雄 (hero) | 形象 (image) | 棺材 (coffin) | 李杰 (Jie Li) |
| 奖金 (bonus) | 时机 (opportunity) | 诡计 (trick) | 张磊 (Lei Zhang) |
| 笑容 (smile) | 首领 (chieftain) | 疾病 (disease) | 李娟 (Juan Li) |
| 智慧 (wisdom) | 光线 (light) | 悲剧 (tragedy) | 王军 (Jun Wang) |
| 冠军 (champion) | 关键 (key) | 谣言 (rumor) | 张艳 (Yan Zhang) |
| 爱情 (love) | 货币 (currency) | 疯子 (madman) | 张涛 (Tao Zhang) |
| 赞叹 (admire) | 爆发 (erupt) | 自杀 (suicide) | |
| 信赖 (trust) | 包围 (encircle) | 屠杀 (massacre) | |
| 体贴 (consider) | 反攻 (counterattack) | 暗杀 (assassin) | |
| 拥抱 (embrace) | 解除 (relieve) | 咒骂 (curse) | |
| 尊重 (respect) | 抵御 (defense) | 侮辱 (insult) | |
| 称赞 (compliment) | 旁观 (observe) | 贪污 (corrupt) | |
| 鼓励 (encourage) | 出兵 (charge) | 虐待 (abuse) | |
| 爱慕 (adore) | 遗留 (remain) | 毁灭 (ruin) | |
| 信任 (believe) | 降临 (befall) | 欺骗 (deceive) | |
| 欢呼 (acclaim) | 摇摆 (sway) | 恐惧 (fear) | |
| 求婚 (propose) | 击败 (defeat) | 背叛 (betray) | |
| 惊喜 (surprise) | 搏斗 (fight) | 怨恨 (hate) | |
| 庆祝 (celebrate) | 降落 (landing) | 灭亡 (perish) | |
| 表扬 (praise) | 告诫 (exhort) | 恐吓 (threaten) | |
| 敬佩 (esteem) | 审查 (censor) | 窒息 (stifle) | |
| 奖励 (award) | 害羞 (shy) | 绑架 (kidnap) | |
| 祝福 (bless) | 脱离 (secede) | 嫉妒 (envy) | |
| 微笑 (smile) | 应酬 (handle) | 出卖 (sell out) | |
| 自信 (assured) | 打败 (defeat) | 恶化 (deteriorate) | |
| 恋爱 (in-love) | 进攻 (attack) | 厌恶 (detest) | |
| 欢快 (bright) | 简易 (simple) | 下贱 (base-bred) | |
| 永恒 (eternal) | 平滑 (smooth) | 卑劣 (despicable) | |
| 乐观 (optimistic) | 高速 (high-speed) | 凶狠 (cruel) | |

| | | |
|---|---|---|
| 满意 (approving) | 正规 (normal) | 可耻 (disgraceful) |
| 杰出 (outstanding) | 详尽 (detailed) | 窝囊 (timid) |
| 深情（affectionateness） | 成年（adult） | 死刑 (death penalty) |
| 真心 (sincerity) | 地位 (status) | 地狱 (hell) |
| 宝石 (jewel) | 首相 (premier) | 恶魔 (demon) |
| 母亲 (mother) | 银行 (bank) | 叛徒 (traitor) |
| 朋友 (friend) | 规律 (rule ) | 凶手 (murderer) |
| 信心 (confidence) | 睡眠 (sleep) | 罪犯 (criminal) |
| 才华 (talent) | 皇帝 (emperor) | 魔鬼 (devil) |
| 友谊 (friendship) | 人情 (relationship) | 罪恶 (evil) |
| 鲜花 (flower) | 军人 (soldier) | 恶意 (venom) |
| 美德 (virtue) | 体操 (gymnastics) | 小偷 (thief) |
| 情侣 (couple) | 权力 (power) | 耻辱 (humiliation) |
| 成就 (achievement) | 人民 (people) | 灾害 (calamity) |
| 天堂 (heaven) | 眼睛 (eye) | 火灾 (conflagration) |
| 美女 (beauty) | 都市 (city) | 奴才 (slave) |
| 英雄 (hero) | 形象 (image) | 棺材 (coffin) |

**Table S2.** Regression model details for external validity.

| | | BDI | | SLA | | PA | | NA | | POS | | NEG | | NET | |
|---|---|---|---|---|---|---|---|---|---|---|---|---|---|---|---|
| Self-reported scores | A | 0.11 | | -0.07 | | -0.00 | | 0.06 | | 0.10 | | 0.19 | | -0.07 | |
| | C | -0.23* | | $0.20^{+}$ | | 0.15 | | -0.26* | | -0.15 | | 0.21 | | 0.06 | |
| | N | 0.38* | | -0.14 | | -0.05 | | 0.43* | | 0.08 | | 0.04 | | -0.32* | |
| | O | 0.20* | | 0.02 | | 0.32* | | 0.13 | | -0.02 | | -0.27 | | -0.22 | |
| | E | -0.25* | | 0.15 | | 0.29* | | -0.12 | | 0.35* | | 0.05 | | 0.19 | |
| EEG-predicted scores | A | | 0.13 | | 0.04 | | -0.08 | | 0.68 | | -0.13 | | -0.10 | | 0.37* |
| | C | | -0.10 | | 0.07 | | 0.14 | | -0.72 | | 0.08 | | -0.04 | | -0.07 |
| | N | | 0.23* | | -0.01 | | -0.11 | | 0.35* | | 0.13 | | -0.29* | | -0.08 |
| | O | | 0.17 | | 0.10 | | 0.17+ | | -0.43 | | 0.04 | | -0.03 | | -0.26* |
| | E | | -0.18 | | 0.12 | | 0.23+ | | 0.14 | | 0.13 | | 0.12 | | 0.29* |
| R | | 0.57* | 0.34* | 0.33* | 0.20 | 0.56* | 0.40* | 0.60* | 0.32* | 0.34 | 0.22 | 0.32 | 0.34 | $0.42^{+}$ | 0.46* |
| $R^2$ | | 0.33* | 0.12* | 0.11* | 0.04 | 0.32* | 0.15* | 0.36* | 0.10* | 0.11 | 0.05 | 0.10 | 0.12 | $0.17^{+}$ | 0.21* |

+: *p* ~~P~~< .1, *: ~~P~~*p* < .05.

BDI is the self-reported score of Beck Depression Inventory, SLAS is the self-reported score of Satisfaction with Life Scale; PA and NA are self-reported scores of negative and positive affects; POS, NEG, NET are participants' ratings of the valence of neutral,negative and positive video clips.

**Table S3.** The video clips list used in the present study.

| Clip Number | Source Film | Targeted Emotion | Duration （s） | Language |
|---|---|---|---|---|
| 1 | Tokyo Trial | Angry | 81 | Japanese |
| 2 | Nanking | Angry | 63 | English |
| 3 | City of Life and Death | Angry | 73 | Chinese |
| 4 | Trainspotting | Disgust | 78 | English |
| 5 | Indiana Jones and the Last Crusade | Disgust | 69 | English |
| 6 | Hellraiser | Disgust | 90 | English |
| 7 | The Shining | Fear | 56 | English |
| 8 | The Shining | Fear | 60 | English |
| 9 | The Exorcist | Fear | 105 | English |
| 10 | In Bruges | Sadness | 45 | English |
| 11 | Departures | Sadness | 60 | Japanese |
| 12 | Gangs of New York | Sadness | 81 | English |
| 13 | Blue 1 | Neutral | 35 | English |
| 14 | Blue 2 | Neutral | 44 | English |
| 15 | Blue 3 | Neutral | 38 | English |
| 16 | The Lover | Neutral | 43 | English |
| 17 | Modern Times | Amusement | 55 | English |
| 18 | Minions | Amusement | 69 | English |
| 19 | Mr. Bean | Amusement | 73 | English |
| 20 | Forrest Gump | Inspiration | 129 | English |
| 21 | The Theory of Everything | Inspiration | 77 | English |
| 22 | The Shawshank Redemption | Inspiration | 83 | English |

| | | | | |
|---|---|---|---|---|
| 23 | My Neighbor Totoro | Joy | 34 | Japanese |
| 24 | Night at the Museum III | Joy | 37 | English |
| 25 | Harry Potter I | Joy | 67 | English |
| 26 | The Pursuit of Happiness | Tenderness | 63 | English |
| 27 | Juno | Tenderness | 83 | English |
| 28 | Sex and the City II | Tenderness | 77 | English |

Notes: To avoid inducing other negative emotion, two angry videos (Rape of Nanking,  City of Life and Death) were re-edited based on the same rule of angry clips from Chinese Emotional Film Clips.
